# UV damage induces G3BP1-dependent stress granule formation that is not driven by translation arrest via mTOR inhibition

**DOI:** 10.1101/2020.04.29.068585

**Authors:** Shan Ying, Denys A. Khaperskyy

**Affiliations:** Department of Microbiology and Immunology, Dalhousie University, Halifax, NS, B3H 4R2, Canada

**Keywords:** stress granule, UVC, GCN2, G3BP1, mTOR

## Abstract

Translation arrest is a part of the cellular stress response that decreases energy consumption and enables rapid reprioritisation of gene expression. Often translation arrest leads to condensation of untranslated messenger ribonucleoproteins (mRNPs) into stress granules (SGs). Studies into mechanisms of SG formation and functions are complicated because various types of stress cause formation of SGs with different properties and composition. In this work we focused on the mechanism of SG formation triggered by UV damage. We demonstrate that UV-induced inhibition of translation does not cause dissociation of the 48S preinitiation complexes. The catalytic activity of the general control non-derepressible 2 (GCN2) kinase contributes to UV-induced SG formation, which is independent of the GCN2-mediated phosphorylation of the eukaryotic translation initiation factor 2α. Like many other types of SGs, condensation of UV-induced granules specifically requires the Ras-GTPase-Activating Protein SH3-Domain-Binding Protein 1 (G3BP1). Our work reveals that in UV-treated cells the mechanisms of translation arrest and SG formation may be unlinked, resulting in condensation of ribonucleoproteins that do not represent the major type of polysome-free preinitiation complexes that accumulate in the cytoplasm.

## INTRODUCTION

Inhibition of translation initiation in response to various types of stress leads to ribosome runoff from messenger RNAs (mRNAs) and condensation of untranslated mRNA-protein complexes (mRNPs) into large foci called stress granules (SGs) (1). SGs are cytoplasmic phase-separated organelles that accumulate polysome-free mRNPs and dozens of proteins and other molecules that are held together by multiple weak RNA-protein and protein-protein interactions (1–3). Assembly of SGs is driven by the SG-nucleating proteins that include the Ras-GTPase-Activating Protein SH3-Domain-Binding Proteins 1 and 2 (G3BP1/2), T-cell internal antigen 1 (TIA-1), and T-cell internal antigen related (TIAR), which are often used as nearly universal SG markers. SGs are believed to play a role in regulating stress responses, however the molecular functions of SG formation remain poorly understood (2, 4).

In some cases aberrant SG dynamics can contribute to neurodegeneration, because phase separation facilitates formation of stable cytotoxic aggregates of mutant proteins linked to neurodegenerative diseases (reviewed in (5–7)). However, transient translation arrest and SG formation are most often discussed in the context of the pro-survival integrated stress response (ISR) program triggered by phosphorylation of eukaryotic initiation factor 2α (eIF2α) on serine-51 by one of the four kinases activated by different types of stress (2, 8, 9). The heme-regulated inhibitor (HRI) kinase is activated by oxidative stress (10); the general control non-derepressible 2 (GCN2) is activated by amino acid starvation (11) or UV damage (12); the double-stranded RNA (dsRNA)-activated protein kinase (PKR) is activated by viral dsRNA replication intermediates (13); and the PKR-like endoplasmic reticulum kinase (PERK) is activated by endoplasmic reticulum stress (14). When eIF2α is phosphorylated, it stably binds eIF2B and prevents it from mediating GDP to GTP exchange that is required for generation of the translation initiation competent eIF2-GTP-Met-tRNA^Met^ ternary complex (15). This inhibits translation initiation downstream of the assembly of the 48S translation preinitiation complex that includes the eIF4F complex (consisting of eIF4E cap binding protein, eIF4G scaffolding subunit, and eIF4A RNA helicase) bound to the mRNA 5’ m^7^GTP cap and the 43S small ribosomal subunit (15). Consequently, the 48S pre-initiation complexes become the major constituents of SGs that form (8).

The phospho-eIF2α dependent translation arrest and SG formation can be blocked by pharmacological inhibition or interference with the expression of specific eIF2α kinases (16), as well as by genetic replacement of the wild-type (WT) eIF2α gene with an unphosphorylatable S51A mutant (17). Recently, a small molecule ISR inhibitor (ISRIB) was developed (18), which does not interfere with eIF2α phosphorylation, but instead blocks its effect of translation initiation by facilitating eIF2B-mediated GDP to GTP exchange (19). When SG formation is dependent on eIF2α phosphorylation, for example in response to treatments with sodium arsenite (As, induces oxidative stress and HRI activation) or thapsigargin (Tg, induces endoplasmic reticulum stress and PERK activation), it is strongly inhibited by ISRIB (19).

SG formation can also be phospho-eIF2α independent. For example, oxidative stress caused by hydrogen peroxide (H_2_O_2_) or sodium selenite (Se) induce SGs through a 4E binding protein (4E-BP) dependent mechanism via inhibition of mechanistic target of rapamycin (mTOR) (20, 21). Under conditions that favour growth and proliferation of cells, mTOR phosphorylates 4E-BP on multiple serine/threonine residues (22, 23). Hyperphosphorylated 4E-BP cannot bind eIF4E, allowing for eIF4E-eIF4G complex formation and assembly of eIF4F. Inhibition of mTOR leads to rapid 4E-BP dephosphorylation and binding and sequestration of eIF4E away from eIF4G. Under these conditions, 48S preinitiation complexes cannot form and translation initiation is inhibited (24). SGs that form via 4E-BP dependent mechanism lack subunits of eIF3 (e.g. eIF3B) because eIF4F complex formation is required for recruitment of eIF3 to the mRNA (20).

One of the stress stimuli that induce SG formation is exposure to UV light. UV exposure is ubiquitous for all living organisms exposed to sunlight. Although high energy UVC (<290 nm wavelength) radiation from the sun is blocked by the ozone layer in the atmosphere, the most dangerous spectrum of UV light that reaches the surface and contributes to the development of skin cancer, UVB (290-320 nm), causes the same types of DNA damage as UVC (25). Accordingly, many studies analysing the effects of UV light on living cells, including our present study, utilize standard 254 nm UVC bulbs. Historically, UV radiation-induced DNA damage and cell cycle arrest has been studied extensively because of their relationship to carcinogenesis (26). By comparison, UV-induced translation arrest, SG formation, and their roles in cell survival following UV damage remain poorly understood. Several studies examining UV-induced SGs revealed that despite causing robust activation of GCN2 and GCN2-mediated eIF2α phosphorylation, UV-induced SG formation does not depend on phospho-eIF2α (16, 27). The UV-induced granules are true SGs because their formation can be inhibited by cycloheximide (CHX) and they accumulate G3BP1/2, TIA-1, TIAR, and FMRP. However, like SGs caused by H_2_O_2_, UV-induced SGs do not accumulate eIF4G and eIF3B, and poorly recruit PABP1 and poly(A) RNA (16, 27). Whether the mechanism of SG formation in response to UV damage is also similar to that of H_2_O_2_ or Se-induced SGs and involves mTOR inhibition has not been investigated.

In this work, we systematically examined translation arrest and SG formation in human U2OS osteosarcoma cells in response to UV light. We demonstrate that the GCN2-mediated eIF2α phosphorylation is responsible for some but not all the UV-induced translation inhibition. By contrast, we did not reveal any contribution of mTOR inhibition or disassembly of eIF4F complex to the translation arrest following UV exposure. Using Me^7^GTP-agarose pulldown assay and co-immunoprecipitation with eIF3B-specific antibody we demonstrate that the bulk of 48S preinitiation complexes remains intact even when translation is strongly inhibited by UV light. Our studies reveal that UV damage triggers a novel mechanism of SG formation that relies neither on eIF2α phosphorylation nor mTOR inhibition. UV-induced SG condensation is driven by G3BP1, it is enhanced by the catalytic activity of GCN2, but recruits only a small fraction of untranslated mRNPs that lose their association with eIF4G and eIF3B.

## MATERIAL AND METHODS

### Cells and treatments

U2OS human osteosarcoma cells were purchased from American Type Culture Collection (ATCC, Manassas, VA, USA). The eIF2α knock-in mouse embryonic fibroblasts (wild-type and S51A mutant) were a kind gift from Dr. Randal Kaufman (Sanford Burnham Prebys Medical Discovery Institute, La Jolla, CA, USA) (17). Unless specified otherwise, reagents were purchased from Thermo Fisher Scientific (Waltham, MA, USA). All cell lines were cultured in Dulbecco’s modified Eagle’s medium (DMEM) supplemented with 10% fetal bovine serum and 2 mM L-glutamine at 37□C in 5% CO2 atmosphere.

For UV light exposure, cells were grown to 65-85% confluency, media was removed, monolayers washed briefly with phosphate buffered saline (PBS) and exposed to 10 or 20 mJ/cm^2^ UV light (254 nm, UVC) in the HL-2000 Hybrilinker chamber (UVP), promptly overlaid with fresh warm media and returned to 37□C incubator.

For amino acid deprivation experiments, the growth medium was replaced with Hanks’ balanced salt solution (HBSS) supplemented with 10% dialysed fetal bovine serum (Wisent Inc., St-Bruno, QC, Canada). For ISRIB or the GCN2 inhibitor treatments, the growth medium was replaced with fresh media containing 200 nM ISRIB (MilliporeSigma, Burlington, MA, USA) or the indicated concentrations of A-92, a.k.a. GCN2-IN-1 (MedChemExpress, Monmouth Junction, NJ, USA). Other treatments were performed by direct addition of 1/100 volume of stocks pre-diluted in media. Final concentrations were as follows: 250 nM Torin-1 (TOCRIS, Oakville, ON, Canada); 250 or 500 µM sodium arsenite; 1 mM sodium selenite; 1 mM hydrogen peroxide (all MilliporeSigma, Burlington, MA, USA).

### Generation of inducible cell line expressing USP10 N-terminal 40 amino acid peptide

Generation of cell lines harbouring doxycycline-inducible protein expression constructs using lentiviral vectors based on the pTRIPZ plasmid (Thermo Fisher Scientific) was previously described in (28). To enable tight control of USP10 N-terminal 40 amino acid peptide expression it was cloned as an EGFP fusion protein into pTRIPZ vector downstream of doxycycline-inducible promoter between AgeI and MluI sites, completely substituting turboRFP and shRNA cassette (MluI site was destroyed during cloning), producing pTRIPZ-EGFP-USP10(1-40) vector (full sequence is available upon request). A companion control plasmid pTRIPZ-EGFP containing EGFP without USP10 peptide fused at the c-terminus was constructed at the same time. U2OS cells were transduced with lentiviruses generated with these vectors at MOI 1.0 and stably transduced cells were selected with 1 µg/ml Puromycin for 48 h. EGFP and EGFP-USP10(1-40) fusion protein expression was induced in resistant cells for 24 h with 0.5 µg/ml doxycycline and EGFP-positive cells were sorted at the Dalhousie University Flow Cytometry Core facility. Sorted cells were propagated for 2 passages without doxycycline and re-tested for doxycycline-regulated EGFP and EGFP-USP10(1-40) fusion protein expression.

### Gene silencing

For GCN2 silencing, U2OS cells were transfected with Ambion Silencer Select Pre-designed siRNAs (s54067, siGCN2-1; or s54069, siGCN2-2) using Lipofectamine RNAiMAX according to manufacturer protocol (reverse transfection in 12-well cluster dishes) and treated/analysed 48 h post-transfection. For non-targeting siRNA control, cells were transfected with Ambion Silencer Select Negative Control #2 (Cat. 4390846).

The lentiCRISPR-v2 plasmids encoding guide RNAs targeting human G3BP1 and G3BP2 genes were a kind gift from Dr. Adrianne Weeks (Dalhousie University, Halifax, NS, Canada). Guide RNA insert sequences: G3BP1 5’-AA GCC TAG TCC CCT GCT GGT CGG; G3BP2 5’-TG GCC ATA AAC AGC TTC CTG GGG. U2OS cells were transduced with lentiviruses generated with these vectors at MOI 1.0 and stably transduced cells were selected with 1 µg/ml Puromycin for 48 h. Resistant cells were seeded onto 12-well cluster dishes and used in experiments 48 h post-seeding (4 days post-transduction with lentiviruses).

### Immunofluorescence staining

Cell fixation and immunofluorescence staining were performed according to procedure described in (29). Briefly, cells grown on 18-mm round coverslips were fixed with 4% paraformaldehyde in PBS for 15 min at ambient temperature and permeabilized with cold methanol for 10 min. After 1-h blocking with 5% bovine serum albumin (BSA, BioShop, Burlington, ON, Canada) in PBS, staining was performed overnight at +4□C with antibodies to the following targets: eIF3B (Rabbit, Bethyl Labs., A301-761A); eIF4G (Rabbit, Cell Signaling, #2498); FMRP (Rabbit, Cell Signaling, #7104); G3BP1 (Mouse, BD Transduction, 611126); TIA-1 (Goat, Santa Cruz, sc-1751); TIAR (Rabbit, Cell Signaling, #8509). Alexa Fluor conjugated secondary antibodies: donkey anti-goat IgG AF488 (Invitrogen, A11055); donkey anti-rabbit IgG AF488 (Invitrogen, A21206); donkey anti-mouse IgG AF555 (Invitrogen, A21202); donkey anti-rabbit IgG AF555 (Invitrogen, A31572); donkey anti-goat IgG AF647 (Invitrogen, A32839); or goat anti-rabbit IgG AF647 (Invitrogen, A21245). Where indicated, nuclei were stained with Hoechst 33342 dye (Invitrogen, H3570). Slides were mounted with ProLong Diamond Antifade Mountant (Invitrogen, P36970) and imaged using Zeiss AxioImager Z2 fluorescence microscope and Zeiss ZEN 2011 software. Quantification of stress granule-positive cells was performed by counting the number of cells with at least two discrete cytoplasmic foci co-stained with two markers from at least 3 randomly selected fields of view, analysing >100 cells per treatment in each replicate.

### Western blotting

Whole cell lysates were prepared by direct lysis of PBS-washed cell monolayers with 1x Laemmli sample buffer (50 mM Tris-HCl pH 6.8, 10% glycerol, 2% SDS, 100 mM DTT, 0.005% Bromophenol blue). To preserve phosphorylation status of proteins, following 5-minute agitation at ambient temperature, lysates were immediately placed on ice, homogenized by passing through a 21-gauge needle, and stored at −20□C. Aliquots of lysates thawed on ice were incubated at 95□C for 3 min, cooled on ice, separated using denaturing polyacrylamide gel electrophoresis, transferred onto PVDF membranes using Trans Blot Turbo Transfer System with RTA Transfer Packs (Bio-Rad Laboratories, Hercules, CA, USA) according to manufacturer protocol and analysed by immunoblotting using antibody-specific protocols. Antibodies to the following targets were used: 4E-BP (Rabbit, Cell Signaling, #9644); β-actin (HRP-conjugated, mouse, Santa Cruz, sc-47778); eIF3A (Rabbit, Cell Signaling, #3411); eIF3B (Rabbit, Bethyl Labs., A301-761A); eIF4E (Rabbit, Cell Signaling, #2067); eIF4G (Rabbit, Cell Signaling, #2498); G3BP1 (Mouse, BD Transduction, 611126); G3BP2 (Rabbit, Bethyl Labs, A302-040A); GCN2 (Rabbit, Cell Signaling, #3302); GFP (Rabbit, Cell Signaling, #2956); LC3B (Mouse, Santa Cruz, sc-376404); phospho-T37/T46-4E-BP (Rabbit, Cell Signaling, #2855); phospho-S51-eIF2α (Rabbit, Cell Signaling, #3398); phospho-T899-GCN2 (Rabbit, Epitomics, 2425-1); phospho-S235/S236-S6 (Rabbit, Cell Signaling, #2211); phospho-T389-S6 kinase (Rabbit, Cell Signaling, #9234). For band visualization, HRP-conjugated anti-rabbit IgG (Goat, Cell Signaling, #7074) or anti-mouse IgG (Horse, Cell Signaling, #7076) were used with Clarity Western ECL Substrate on the ChemiDoc Touch Imaging Sysytem (Bio-Rad Laboratories, Hercules, CA, USA).

### Ribopuromycylation assay

Puromycin incorporation assay was performed as described in (30) with the following modifications: puromycin was added to the media at the final concentration of 10 µg/ml for 10 min. Cells were washed with PBS and the whole cell lysate preparation and western blotting analysis were done as described above. For electrophoresis, samples were loaded onto Mini-PROTEAN TGX Pre-cast Stain-Free gels (5-15%, Bio-Rad Laboratories, Hercules, CA, USA) and total protein was visualised post-transfer to PVDF membranes on ChemiDoc Touch Imaging System. Puromycin incorporation into nascent polypeptides was visualised using anti-puromycin antibody (Mouse, MilliporeSigma, MABE343), quantified using Bio-Rad Image Lab 5.2.1 software and values normalised to StainFree signal for each lane.

### m^7^GTP-agarose pulldown assay

m^7^GTP cap pulldown assay was performed as described in (21), except we used γ-Aminophenyl-m7GTP agarose (AC-155, Jena Biosciences GmbH, Jena, Germany). Briefly, cells grown in 10-cm dishes were lysed with buffer containing 50 mM Tris-HCl pH 7.4, 100 mM NaCl, 1 mM EDTA, 0.5% Igepal (NP-40 substitute), and protease/phosphatase inhibitor cocktail (Cell Signaling, #5872). To decrease non-specific binding to beads, lysates were pre-cleared using blank agarose (AC-001, Jena Biosciences GmbH, Jena, Germany) for 15 min at +4□C prior to γ-Aminophenyl-m7GTP agarose pulldown. Binding was performed for 2 h at +4□C with rotation, followed by 3 washes with lysis buffer and elution with 1x Laemmli sample buffer for 10 min at +65□C.

### Co-immunoprecipitation using anti-eIF3B antibody

Cells grown in 10-cm dishes were lysed with ice-cold buffer containing 50 mM Tris-HCl pH 7.6, 140 mM NaCl, 1.5 mM MgCl_2_, 0.5% Igepal (NP-40 substitute), and protease/phosphatase inhibitor cocktail (Cell Signaling, #5872). After 5 min, cells were scraped and incubated for 5 min at +4□C with rotation. Lysates were clarified for 5 min at 10,000 × g at +4□C and protein concentration was determined using DC Protein Assay (Bio-Rad Laboratories, Hercules, CA, USA). For immunoprecipitation, 1 µg of rabbit anti-eIF3B antibody (Bethyl Labs., A301-761A) was added to each lysate containing 0.5 mg of protein in 750 µl volume and incubated for 1 h at +4□C with rotation. Then, 75 µl of protein G magnetic bead suspension (S1430, NEB) pre-equilibrated in lysis buffer was added to each tube and incubation continued for additional 1 h at +4□C. Beads were concentrated using magnetic rack, supernatant removed, and beads were washed 3 times with 500 µl of lysis buffer using magnetic rack according to manufacturer protocol. Co-immunoprecipitated proteins were eluted from the beads using 150 µl of 1x Laemmli sample buffer for 5 min at +75□C and analysed by western blotting as described above except the HRP-conjugated mouse anti-rabbit native IgG secondary antibody (Cell Signaling, #5127) was used to prevent visualisation of heavy and light chains of co-eluted antibody.

### Statistical analysis

All numerical values are plotted as means calculated from 3 independent biological replicates (separate experiments performed on different days), the error bars represent standard deviations. Statistical analysis of each data set is described in figure legends and was performed using GraphPad Prism 8 software. Asterisks denote p values as follows: * p < 0.05; ** p < 0.01; *** p < 0.001.

## RESULTS

### UV-induced SG formation is not associated with inhibition of mTOR signaling

To investigate the role of mTOR signaling in UV-induced translation arrest, we treated U2OS cells with 10 or 20 mJ/cm^2^ UVC light and analysed phosphorylation of mTOR targets 4E-BP and S6K by western blotting (Fig. 1A) and protein synthesis rates using a ribopuromycylation assay (Fig. 1B) at 2 h post-UV treatment. We chose this time point for all analyses because even though SG formation is easily detectable at 1 h post-UV exposure (16), we previously determined that SG formation peaks around 2 h post-UV exposure in these cells (31). For a positive control, we used the catalytic inhibitor of mTOR Torin-1 (32). As expected, Torin-1 treatment decreased phosphorylation of 4E-BP and S6K to nearly undetectable levels in both UV-treated and untreated cells. It also induced autophagy, as measured by increased LC3B-II levels. By contrast, no changes in S6K phosphorylation or LC3B-II levels were observed following UV treatment. Interestingly, 4E-BP became hyperphosphorylated following UV exposure, as indicated by the appearance of phospho-4E-BP bands with reduced electrophoretic mobility (Fig. 1A, compare lane 1 to 3 and 5). The UV-induced GCN2 autophosphorylation and the GCN2-mediated eIF2α phosphorylation were not affected by mTOR inhibition, indicating that under conditions tested, mTOR is not required for GCN2 activation and signalling. The ribopuromycylation assay showed that mTOR inhibition had an additive effect to the UV-induced translation arrest (Fig. 1B), with the magnitude of translation arrest induced by UV treatment alone not corresponding to the levels of mTOR inhibition (Fig. 1A and 1B, compare lane 2 to lanes 3 and 5).

**Figure 1.**
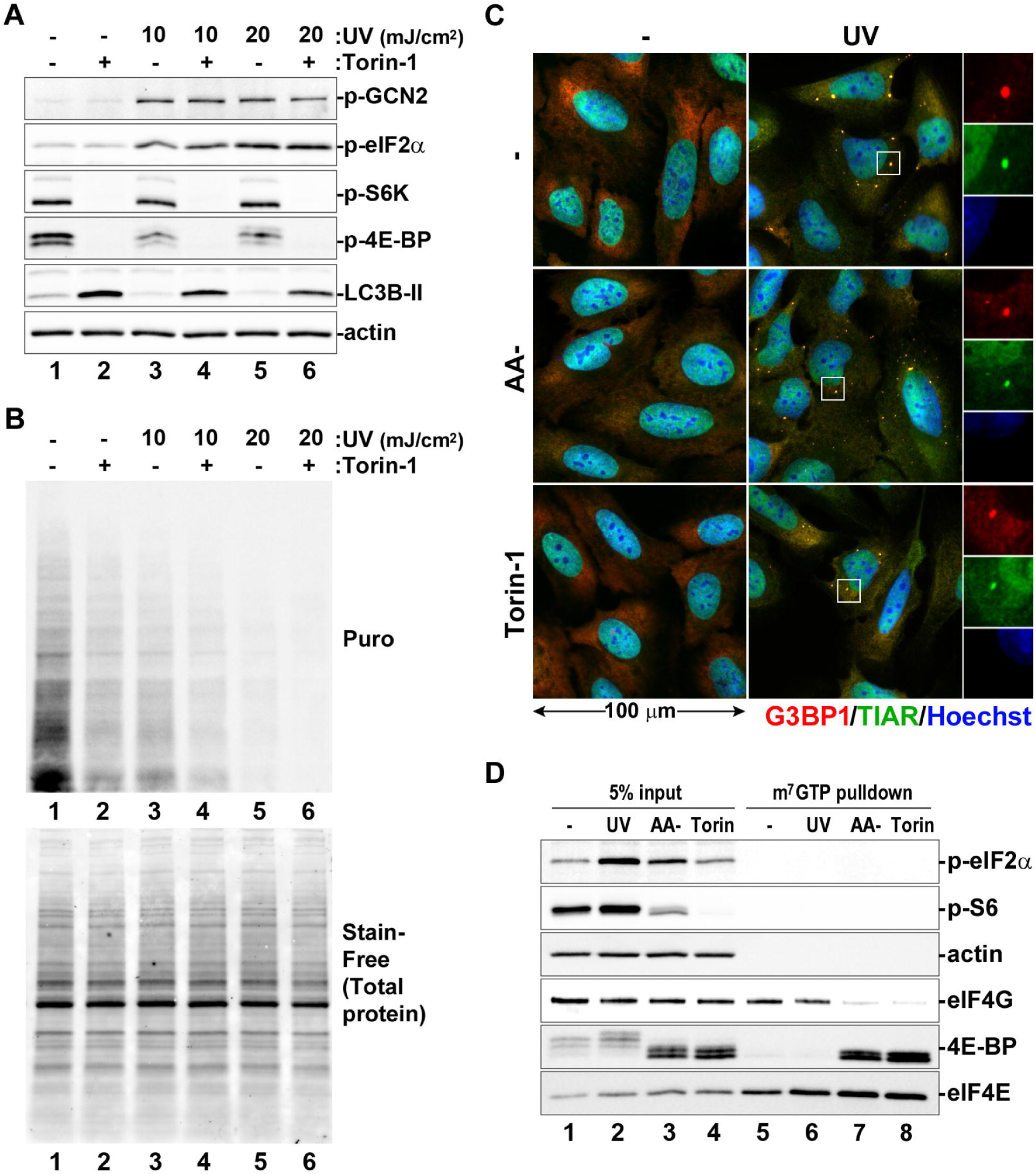
Translation arrest and SG formation in response to UV damage is independent of mTOR inhibition. (A and B) U2OS cells were exposed to the indicated doses of UV light and incubated for 2 h with or without Torin-1 prior to analysis. (A) Phosphorylation of GCN2, eIF2α, mTOR substrates 4E-BP and S6K, as well as lipidation of LC3B (LC3B-II) were analysed by western blotting. Staining for actin was used as loading control. (B) Protein synthesis rates were analysed using ribopuromycylation assay and western blotting with anti-puromycin antibody (Puro, top panel). Total protein content in each lane was visualised using Stain-Free reagent (bottom panel). (C) UV-induced SG formation was analysed in control cells or cells incubated in amino acid-free media (AA-) or Torin-1 containing media using immunofluorescence staining for G3BP1 (red) and TIAR (green). Nuclei were stained with Hoechst dye (blue). (D) eIF4E-eIF4G and eIF4E-4E-BP interactions in cells treated with UV light (UV), amino acid-free media (AA-), or Torin-1 were analysed by m7GTP-agarose pulldown assay and western blotting. Phosphorylation status of eIF2α and S6 ribosomal protein in whole cell lysates (5% input) were determined by staining with specific antibodies. Staining for eIF4E was used as positive, and for actin as negative controls for m7GTP pulldown.

Having determined that UV-induced translation arrest is independent of mTOR inhibition, we tested how mTOR inhibition affects UV-induced SG formation. Immunofluorescence microscopy analysis of cells treated with UV light showed that inhibition of mTOR by either Torin-1 or amino acid starvation did not prevent SG formation (Fig. 1C). We used amino acid starvation as another control in this experiment, because, unlike UV light, it is known to cause both the GCN2-dependent eIF2α phosphorylation (33) and mTOR inhibition (34). SGs did not form in cells incubated in amino acid free media without UV exposure, indicating that the eIF2α phosphorylation by GCN2 is not sufficient to drive SG condensation when mTOR is inhibited. Strong inhibition of 4E-BP phosphorylation in Torin-1 treated cells should cause disassembly of eIF4F translation preinitiation complex, and our results show that UV-induced SGs can still form in Torin-1 treated cells (Fig. 1C). This suggests that UV-induced SGs can accumulate untranslated mRNPs that lack eIF4F. To determine if UV damage causes eIF4F complex disassembly that is independent of mTOR inhibition, we compared eIF4F complex formation in UV-treated cells and cells treated with either amino acid free media or Torin-1 using m^7^GTP-agarose pulldown assay (Fig. 1D). As expected, mTOR inhibition by amino acid starvation or Torin-1 caused dephosphorylation of 4E-BP, its association with eIF4E, and dissociation of eIF4E and eIF4G (Fig. 1D, lanes 7 and 8). By contrast, UV treatment did not affect eIF4E-eIF4G interaction and did not increase 4E-BP association with eIF4E (Fig. 1D, lane 6). This indicates that SG condensation in response to UV does not correlate with eIF4F complex disassembly nor does it depend on the bulk of untranslated mRNPs which maintain an intact eIF4F complex (Fig. 1C,D).

### eIF2α phosphorylation is not required for UV-induced SG formation

To confirm that eIF2α phosphorylation is not required for SG formation in response to UV damage, we compared translation arrest and SG formation in mouse embryonic fibroblasts (MEFs) engineered to express either the wild-type eIF2α (MEF[eIF2α-WT]) or the unphosphorylatable S51A mutant (MEF[eIF2α-S51A]) (17). In these experiments we used sodium arsenite as a control, because it is known to induce phospho-eIF2α dependent translation arrest (10). Treatment of MEF[eIF2α-WT] cells with UV light caused increased eIF2α phosphorylation and decreased protein synthesis rates in a dose-dependent manner, as determined by western blotting and ribopuromycylation assays, respectively (Fig. 2A,B; lanes 3 and 4). In these cells, sodium arsenite triggered eIF2α phosphorylation levels and translation arrest comparable to those induced by 20 mJ/cm^2^ UV (Fig. 2A,B; compare lanes 2 and 4). As expected, in MEF[eIF2α-S51A] cells no phospho-eIF2α signal was detected by western blotting (Fig. 2A, lanes 5-8), yet substantial decrease in protein synthesis rates was evident following UV treatment (Fig. 2B, compare lane 5 to lanes 7 and 8), indicating that UV-induced translation arrest is phospho-eIF2α independent. Only ∼20% translation arrest was induced by sodium arsenite in MEF[eIF2α-S51A] cells (Fig. 2B, lane 6), confirming that arsenite inhibits translation via a phospho-eIF2α dependent mechanism. To gain further insight into the exact contribution of eIF2α phosphorylation to translation inhibition, we carefully quantified puromycin incorporation from 3 independent experiments comparing treatment with sodium arsenite to exposure to 20 mJ/cm^2^ UV. Experiments were performed as shown in Fig. 2A and B, except we introduced treatment with the integrated stress response inhibitor (ISRIB) as an additional control to block effects of eIF2α phosphorylation on translation initiation (18, 19). In MEF[eIF2α-WT] cells, ISRIB increased protein synthesis rates after both arsenite and UV treatment (Fig. 2C). However, even in the presence of ISRIB, UV light caused more than 60% inhibition of translation. By contrast, incubation with ISRIB had no effect on protein synthesis rates in MEF[eIF2α-S51A] cells, as expected, with UV exposure causing over 3 times greater inhibition of translation compared to arsenite (Fig. 2C). Next, we analysed UV-induced SG formation in MEF[eIF2α-S51A] cells using immunofluorescence staining and saw that SGs formed in these cells just as well as in MEF[eIF2α-WT] cells (Fig. 2D). Together, these experiments suggest that despite strong eIF2α phosphorylation, UV-induced SG formation may be largely driven by phospho-eIF2α independent translation arrest.

**Figure 2.**
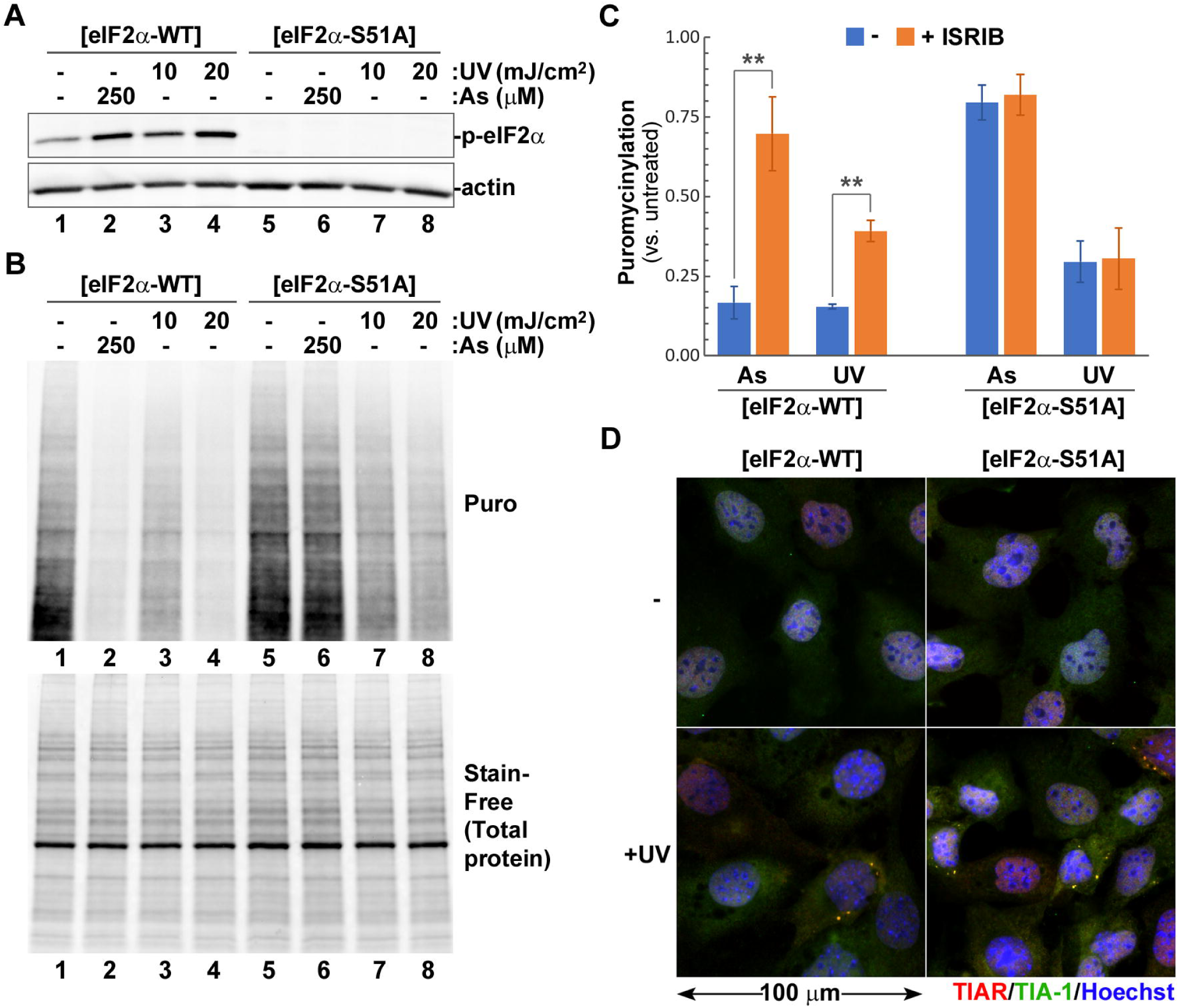
UV damage triggers phospho-eIF2α-independent translation arrest and SG formation. UV-induced translation arrest and SG formation were analysed in mouse embryonic fibroblasts engineered to express either wild-type or S51A mutant eIF2α ([eIF2α-WT] or [eIF2α-S51A], respectively). (A) eIF2α phosphorylation in response to sodium arsenite (As) or UV light (UV) were analysed by western blotting. Staining for actin was used as loading control. (B) Protein synthesis rates were analysed in cells treated with sodium arsenite (As) or UV light (UV) using ribopuromycylation assay and western blotting with anti-puromycin antibody (Puro, top panel). Total protein content in each lane was visualised using Stain-Free reagent (bottom panel). (C) Sodium arsenite (As) and UV-induced translation arrest was quantified from ribopuromycylation assays of cells incubated with or without ISRIB (N = 3). Error bars represent standard deviation; **p < 0.01 (Two-way ANOVA followed by Tukey’s multiple comparisons test). (D) SG formation in UV treated (+UV) and control (-) cells was visualised by staining for TIAR (red) and TIA-1 (green). Nuclei were stained with Hoechst dye (blue).

### GCN2 partially contributes to translation arrest and SG formation in response to UV damage

To examine how GCN2 contributes to UV-induced translation arrest and SG formation, we compared these responses in control U2OS cells, cells transfected with non-targeting siRNA, and cells in which GCN2 expression was silenced using transfection with 2 specific siRNAs (Fig. 3). Western blotting analysis of whole cell lysates at 48 h post-transfection demonstrated that both GCN2-specific siRNAs caused significant depletion of total GCN2 protein compared to untransfected cells or cells transfected with control non-targeting siRNA (Fig. 3A). Importantly, siRNA knock down of GCN2 resulted in diminished eIF2α phosphorylation in response to UV treatment (Fig. 3A, lanes 7 and 8). Ribopuromycylation assay revealed that in cells transfected with one of the two siRNAs (siGCN2-1), the basal rates of protein synthesis were decreased compared to cells transfected with either siGCN2-2 or non-targeting siRNA control (Fig. 3B, compare lane 3 to lanes 2 and 4). This effect, however, was likely caused by unknown off-target effects, as the degree of GCN2 silencing was comparable between two siRNAs (Fig. 3A, compare lanes 3, 4, 7 and 8). When we quantified rates of protein synthesis in control and GCN2-silenced cells following UV treatment, we observed that GCN2 knock down partially restored translation (Fig. 3C). Interestingly, in cells transfected with either GCN2-specific siRNA, the magnitude of UV-induced translation arrest was not affected by ISRIB treatment and was generally comparable to the magnitude of UV-induced translation arrest in control cells treated with ISRIB. This suggests that the difference in translation arrest between control and GCN2 knock down cells can be largely attributed to the effects of GCN2-mediated eIF2α phosphorylation. Next, we analysed UV-induced SG formation in control and GCN2 knock down cells by immunofluorescence staining (Fig. 3D) and quantified SG formation from 3 independent experiments (Fig. 3E). Our analyses revealed that GCN2 silencing decreased SG formation, with siGCN2-1 having more profound effect than siGCN2-2 (Fig. 3D,E). ISRIB treatment had no effect on SG formation, except for a slight decrease in untransfected cells which was statistically significant. Given the profound effects of siGCN2-1 on SG formation and its effects on basal protein synthesis rates in untreated cells, we compared the effects of siGCN2-1 and siGCN2-2 on arsenite-induced SG formation. Sodium arsenite induces SG formation independently from GCN2 through eIF2α phosphorylation by HRI (10), yet siGCN2-1 transfection severely affected SG formation following arsenite treatment (Fig. 3F). This indicates that siGCN2-1 has strong off-target effects that interfere with SG formation in general and cannot be used for analyses of GCN2 effects specifically. Nevertheless, given that siGCN2-2 inhibited SG formation in response to UV but not sodium arsenite, our results suggest that GCN2 contributes to the efficiency of SG formation caused by UV damage.

**Figure 3.**
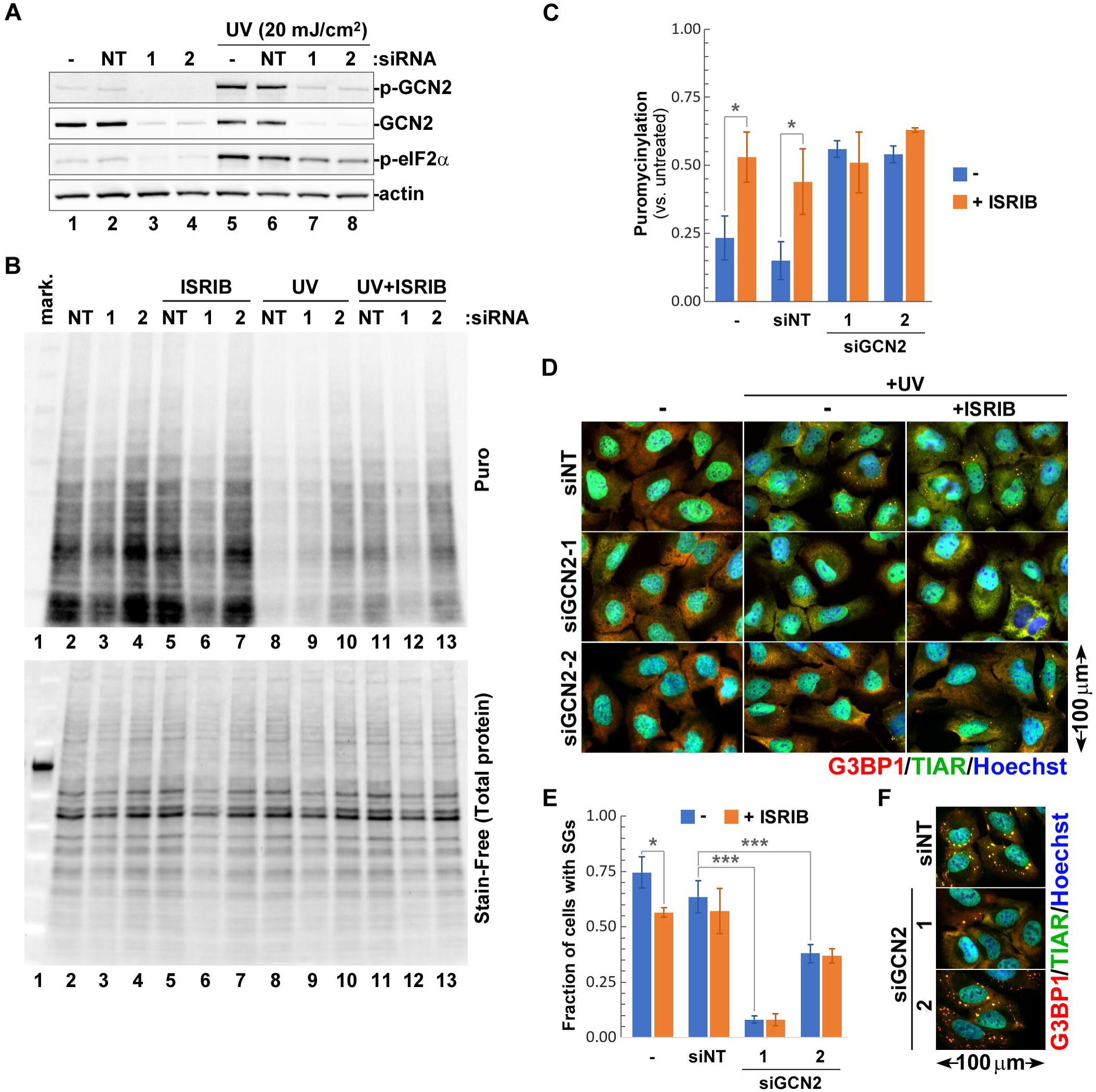
UV-induced SG formation is enhanced by GCN2. U2OS cells were transfected with two different siRNAs targeting GCN2 (1 or 2), non-targeting siRNA (NT), or left untransfected, and were analysed after 48 h for GCN2 expression levels, UV-induced translation arrest and SG formation. (A) Effects of GCN2 silencing on eIF2α phosphorylation in response to UV light (UV) were analysed by western bloting. Efficiency of siRNA-mediated knock down was monitored by staining for total and phosphorylated GCN2. Staining for actin was used as loading control. (B-E) Effects of GCN2 and GCN2-mediated eIF2α phosphorylation on translation arrest and SG formation were analysed in cells incubated with or without ISRIB for 2 h post-UV exposure. (B) Protein synthesis rates were analysed using ribopuromycylation assay and western blotting with anti-puromycin antibody (Puro, top panel). Total protein content in each lane was visualised using Stain-Free reagent (bottom panel). (C) UV-induced translation arrest was quantified from ribopuromycylation assays (N = 3). Error bars represent standard deviation; *p < 0.05 (Two-way ANOVA followed by Tukey’s multiple comparisons test). (D) Immunofluorescence microscopy staining of control or UV-treated (+UV) cells incubated with or without ISRIB for 2 h post-UV exposure. SG formation was analysed by staining for G3BP1 (red) and TIAR (green). Nuclei were stained with Hoechst dye (blue). (E) UV-induced SG formation was quantified from immunofluorescence microscopy staining (N = 3). Error bars represent standard deviation; *p < 0.05; **p < 0.01; ***p < 0.001 (Two-way ANOVA followed by Tukey’s multiple comparisons test). (F) SG formation in cells treated with sodium arsenite was analysed by staining for G3BP1 (red) and TIAR (green). Nuclei were stained with Hoechst dye (blue).

### Catalytic activity of GCN2 enhances UV-induced SG formation

Our analysis of GCN2 contributions to translation arrest and SG formation in response to UV damage using siRNA silencing revealed that GCN2-mediated phosphorylation of eIF2α is responsible for approximately 1/3 of the translation inhibition following UV exposure (Fig. 3C). Interestingly, while eIF2α phosphorylation did not contribute to SG formation, silencing of GCN2 consistently decreased SG formation by at least 30% (Fig. 3E). To determine if GCN2 catalytic activity is important for UV-induced SG formation, we analysed formation of SGs following 20 mJ/cm^2^ UV exposure in cells treated with GCN2 inhibitor A-92 (a.k.a. GCN2-IN-1) and in control DMSO-treated cells (Fig. 4). First, we established the optimal concentration of A-92 by analysing eIF2α phosphorylation in response to UV light by western blotting (Fig. 4A). Unexpectedly, the highest concentration tested (20 µM A-92) caused increased phospho-eIF2α even in untreated cells (Fig. 4A, compare lanes 1 and 2), possibly due to off-target cytotoxicity. For analyses of SG formation, we used 5 mM A-92 treatment because it was sufficient for preventing eIF2α phosphorylation without major effects on the upstream activation and autophosphorylation of GCN2 itself (Fig. 4A, compare lanes 6, 7, and 8). Immunofluorescence staining of cells at 2 h post-UV treatment revealed that compared to DMSO control, A-92 caused formation of slightly smaller SGs in fewer cells (Fig. 4B). Quantification of SG-positive cells showed significant decrease in the presence of A-92, confirming that the catalytic activity of GCN2 is important for SG formation in response to UV damage. Since UV-induced SG formation is phospho-eIF2α independent, our data points to a potential involvement of other yet to be identified GCN2 targets.

**Figure 4.**
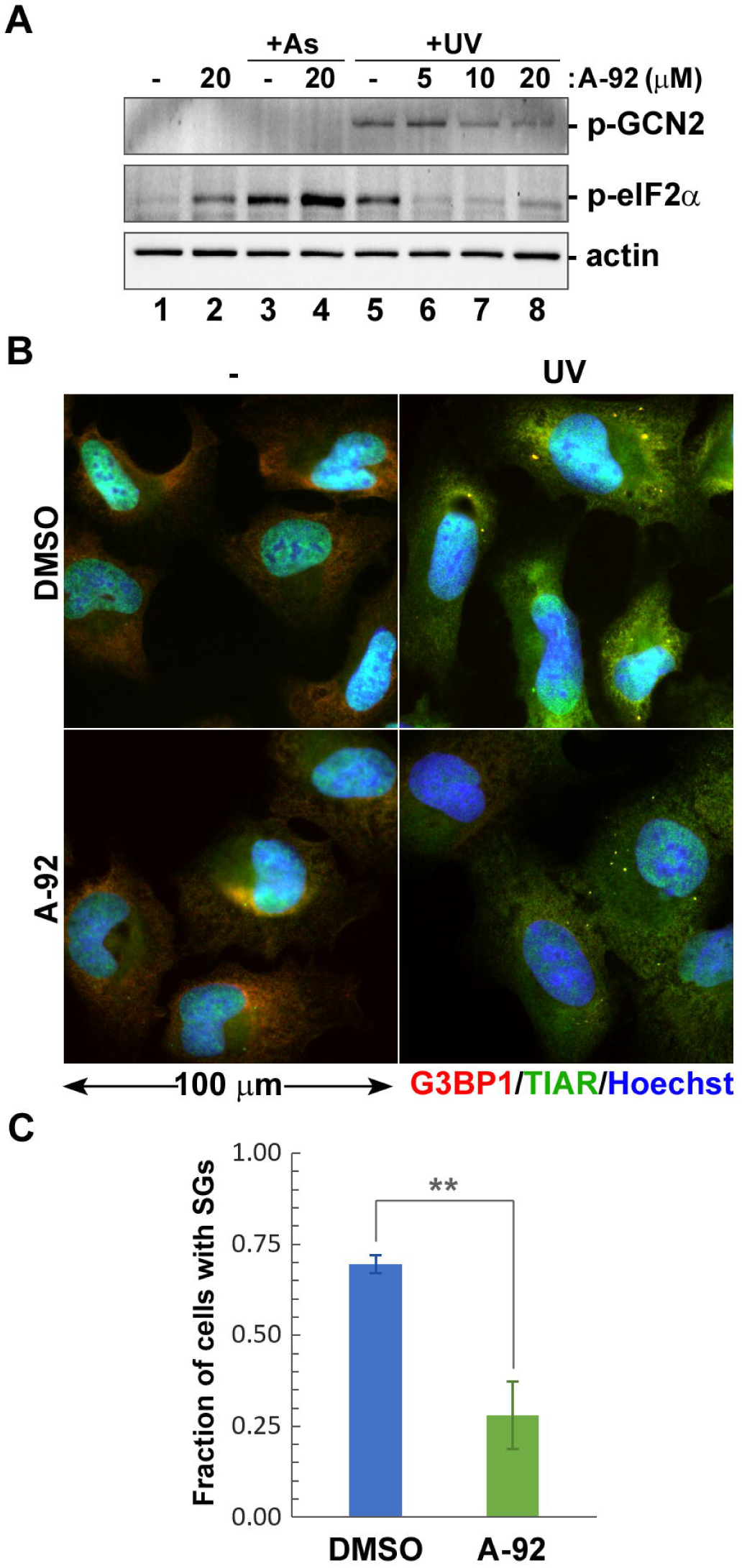
Catalytic activity of GCN2 is important for SG formation. (A) Concentration-dependent effects of A-92 on GCN2 and eIF2α phosphorylation in response to UV light (+UV) were analysed by western blot. Lysates from untreated cells and cells treated with sodium arsenite (+As) were used as controls. Staining for actin was used as loading control. (B) Immunofluorescence microscopy staining of control (-) or UV-treated (UV) cells incubated with 5 µM A-92 or vehicle (DMSO) for 2 h post-exposure. SG formation was visualised by staining for TIAR (red) and TIA-1 (green). Nuclei were stained with Hoechst dye (blue). (C) UV-induced SG formation in control or A-92 treated cells was quantified from immunofluorescence microscopy staining (N = 3). Error bars represent standard deviation; **p < 0.01 (two-tailed Student’s t-test).

### UV-induced SG condensation requires G3BP1 but not G3BP2

Different types of SGs require different nucleating proteins, with G3BP1 and/or G3BP2 being necessary for SG formation in response to variety of stresses, with just a few notable exceptions (35). To test if UV-induced SGs require G3BP1 and/or G3BP2 for their formation, we disrupted expression of these proteins using transduction with lentiviruses encoding CRISPR/Cas-9 cassettes with guide RNAs specific for G3BP1 or G3BP2 and analysed SG formation using immunofluorescence microscopy (Fig. 5A). As controls, we used treatments with sodium arsenite and sodium selenite, which reliably induce robust SG formation in U2OS cells in phospho-eIF2α dependent (arsenite) and independent (selenite) manner (10, 21). For SG detection, we used antibodies to G3BP1, FMRP, and TIA-1 markers. At 4 days following lentiviral transduction, over 80% of cells expressing guide RNAs for G3BP1 (lentiCRISPR-v2-G3BP1) had undetectable levels of this protein, with remaining cells serving as internal controls (Fig. 5A, top row). As expected, both G3BP1-positive and G3BP1-negative cells formed SGs following sodium arsenite treatment, as it was shown that only simultaneous knock out of G3BP1 and G3BP2 can block arsenite-induced SG formation (35). Similarly, G3BP1-negative cells formed SGs following treatment with sodium selenite, as revealed by co-staining with FMRP and TIA-1. By contrast, only G3BP1-positive cells could form UV-induced SGs (Fig. 5A, top row). Unfortunately, we could not detect G3BP2 knockout cells using immunofluorescence, but our western blotting analysis confirmed that G3BP2 was depleted from lentiCRISPR-v2-G3BP2 transduced cells (Fig. 5B). In these cells, formation of arsenite and UV-induced SGs was not affected. Surprisingly, G3BP2 depletion blocked selenite-induced SG formation (Fig. 5A, bottom row). Thus, our analysis reveals that UV-induced SGs require G3BP1, but not G3BP2, for their formation.

**Figure 5.**
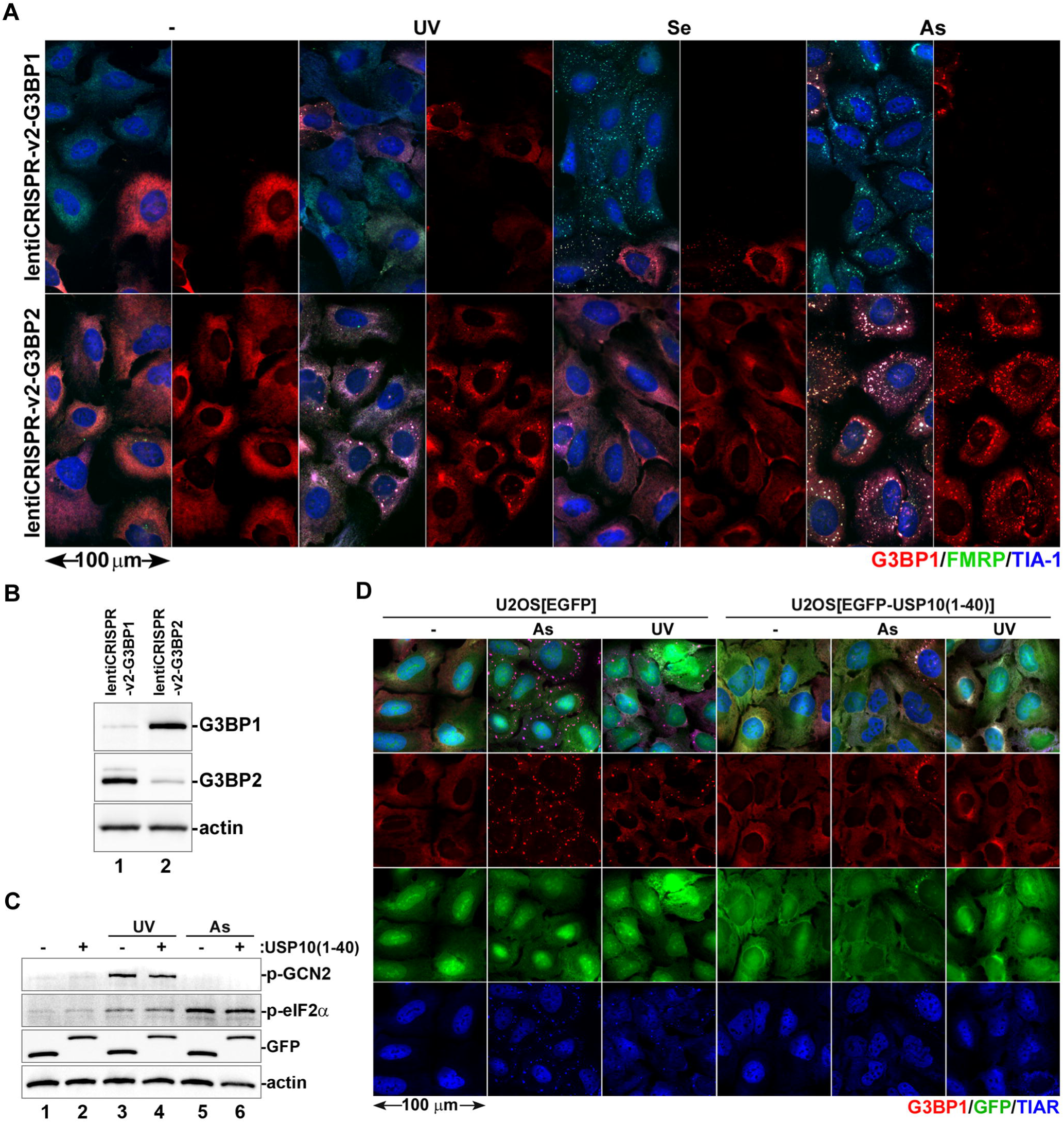
UV-induced SG formation is dependent on G3BP1. (A and B) U2OS cells were transduced with lentivirus vectors bearing CRISPR/Cas9 cassettes targeting G3BP1 (lentiCRISPR-v2-G3BP1) or G3BP2 (lentiCRISPR-v2-G3BP2) and propagated for 4 days. (A) SG formation was analysed by immunofluorescence microscopy in cells lacking G3BP1 (top row) or G3BP2 (bottom row) treated with UV light (UV), sodium selenite (Se), or sodium arsenite (As) and stained for G3BP1 (red), FMRP (green), and TIA-1 (blue). Staining in red channel show few remaining G3BP1-positive cells in the top row that serve as internal controls. (B) Expression of G3BP1 and G3BP2 proteins was analysed by western blotting. Staining for actin was used as loading control. (C and D) Expression of the EGFP-USP10(1-40) fusion protein or EGFP control was induced in U2OS cells stably transduced with lentiviral constructs encoding these proteins. After 48 h, cells were left untreated (-), or were treated with UV light (UV) or sodium arsenite (As). (C) Phosphorylation status of GCN2 and eIF2α post-treatment was analysed by western blotting, as well as the expression of EGFP-USP10(1-40) fusion protein or EGFP control. Staining for actin was used as loading control. (D) SG formation was analysed by immunofluorescence microscopy staining for G3BP1 (red) and TIAR (blue) in these GFP-positive cells (green).

G3BP1-dependent SG condensation requires interaction of its NTF-2-like domain with Caprin-1 protein, and this interaction can be disrupted by the N-terminal 40-amino acid long region of USP10 (35, 36). To test if Caprin-1 dependent mechanism of SG condensation is required for UV-induced SG formation, we generated U2OS cells carrying doxycycline-inducible EGFP fused to the N-terminal 40 amino-acid peptide of USP10 (U2OS[iEGFP-USP10(1-40)]) and the control U2OS[iEGFP] cells. Following induction of EGFP-USP10(1-40) or control EGFP protein expression, we treated these cells with UV or sodium arsenite and analysed GCN2 and eIF2α phosphorylation using western blotting (Fig. 5C) and SG formation using immunofluorescence staining (Fig. 5D). As expected, the eIF2α phosphorylation in response to UV or sodium arsenite was not affected by fusion protein expression (Fig. 5C). By contrast, SG formation in response to both treatments was blocked in EGFP-USP10(1-40) expressing cells (Fig. 5D). Thus, UV-induced SG condensation requires G3BP1 and can be blocked by disrupting interactions of its NTF-2-like domain.

### UV-induced SGs do not accumulate eIF4G and eIF3B

All our results so far suggest that UV damage induces G3BP1-dependent SG formation that does not require mTOR inhibition or eIF2α phosphorylation – two main pathways that can lead to the influx of untranslated mRNPs in the cytoplasm of cells under stress. Previous studies have indicated that UV-induced SGs lack eIF4G and eIF3B (16). These proteins are associated with stalled 48S pre-initiation complexes that accumulate following translation arrest induced by eIF2α phosphorylation (8). To confirm that UV-induced SGs indeed lack eIF4G and eIF3B in our system, we analysed SG composition using immunofluorescence microscopy. As controls, we used treatments with sodium arsenite, hydrogen peroxide, and sodium selenite. The latter two treatments were shown to induce SGs that do not accumulate eIF3B (20, 21). Interestingly, in our system both eIF4G and eIF3B were recruited to some SGs formed in response to hydrogen peroxide and sodium selenite (Fig. 6A,B), which may indicate slight differences in experimental conditions and/or specificity of antibodies used in our study. By contrast, our analysis revealed that UV-induced SGs did not recruit eIF4G or eIF3B in U2OS cells (Fig. 6A,B), indicating that despite the majority of mRNPs maintaining interaction with eIF4G following UV exposure (Fig. 1D), they do not represent the main constituents of SGs in our system. Next, to determine if eIF4G-eIF3B interactions are disrupted after UV treatment of U2OS cells, we conducted co-immunoprecipitation assays using the same anti-eIF3B antibody that we used for immunofluorescence staining in Fig. 6B. Torin-1 served as a positive control, whereas sodium arsenite served as a negative control that does not disrupt integrity of 48S pre-initiation complexes. As expected, in untreated and sodium arsenite treated cells, similar amounts of eIF4G co-immunoprecipitated with eIF3B (Fig. 6C, lanes 5 and 8). In Torin-1 treated cells, eIF4G did not co-immunoprecipitate with eIF3B (Fig. 6C, lane 7). In UV-treated cells, eIF4G-eIF3B interaction was not disrupted and amounts of co-immunoprecipitated eIF4G were similar to those in untreated or sodium arsenite treated cells (Fig. 6C, lane 6). It is important to note that under all conditions tested, eIF3A, another subunit of eIF3 core complex, co-immunoprecipitated efficiently with eIF3B (Fig. 6C). Thus, our data is consistent with the working model where most 48S pre-initiation complexes remain intact after UV damage, similar to those that accumulate in cells treated with sodium arsenite. At the same time, UV-induced SGs form via the G3BP1-dependent mechanism but accumulate only a fraction of untranslated mRNPs that lack eIF4G and eIF3B (Fig. 6D).

**Figure 6.**
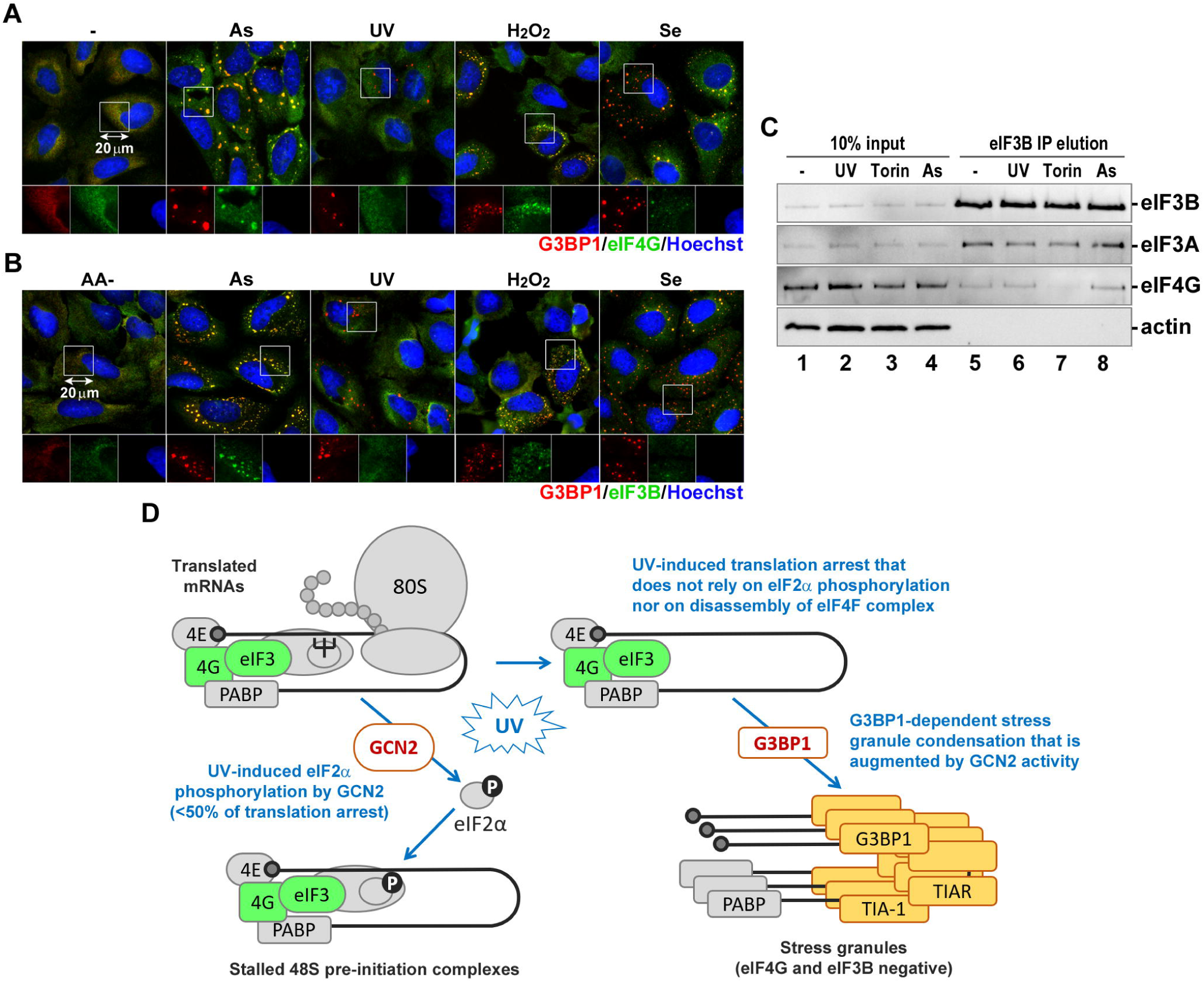
Ribonucleoproteins recruited to UV-induced SGs selectively exclude eIF4G and eIF3B. (A) Cells treated with sodium arsenite (As), UV light (UV), hydrogen peroxide (H_2_O_2_), or sodium selenite (Se) were analysed by immunofluorescence staining for SG markers G3BP1 (red) and eIF4G (green). Nuclei were stained with Hoechst dye (blue). (B) Cells incubated in amino acid-free medium (AA-) or treated with sodium arsenite (As), UV light (UV), hydrogen peroxide (H_2_O_2_), or sodium selenite (Se) were analysed by immunofluorescence staining for SG markers G3BP1 (red) and eIF3B (green). Nuclei were stained with Hoechst dye (blue). (C) Association of eIF3 complex with eIF4G was analysed using co-immunoprecipitation from cytoplasmic lysates of untreated control cells (-) or cells treated with UV light (UV), Torin-1 (Torin), or sodium arsenite (As). Presence of eIF3A and eIF4G proteins in cytoplasmic lysates (10% input) and eIF3B co-immunoprecipitation samples (IP elution) was analysed by western blotting. Staining for eIF3B was used as positive, and for actin as negative controls. (D) Schematic diagram depicting the working model for UV-induced translation arrest and SG formation. GCN2-mediated eIF2α phosphorylation is only partially contributing to overall translation arrest, however most untranslated messenger ribonucleoproteins remain associated with eIF4G and eIF3. SGs form in G3BP1-dependent manner but recruit only a small fraction of untranslated mRNAs and exclude eIF4G and eIF3.

## DISCUSSION

Depending on the type, magnitude, or duration of stress, SG formation was linked with cell survival, apoptosis, regulation of cell cycle or innate immune signaling (27, 37–39). Nevertheless, the exact molecular functions of these condensates remain poorly understood. Studies into mechanisms of SG formation and functions are complicated by the fact that various types of stress trigger formation of SGs that differ significantly in their properties and composition (16). In this work we focused on the mechanisms of translation arrest and SG formation triggered by UV damage. When cells are exposed to various doses of UV, they arrest the cell cycle until the DNA damage induced by UV light can be repaired (40). High dose UV exposure also causes translation arrest and SG formation (12, 16, 27, 41). In cultured mammalian cells, doses of 1 mJ/cm^2^ and above are sufficient for cell cycle arrest (25), however doses as high as 5 mJ/cm^2^ are required for induction of eIF2α phosphorylation and inhibition of protein synthesis and up to 20 mJ/cm^2^ for reliable SG induction at 1-2 h post-exposure (16, 29). Since our analyses were focused specifically on SG formation and similar doses were used by others previously, we decided to use doses of 10-20 mJ/cm^2^ in our experiments.

The “canonical” stress granules form as part of the ISR and their formation is initiated by phosphorylation of eIF2α by one of the four kinases that become activated by different types of stress. The classical example of “canonical” SGs are those induced by treatment with sodium arsenite – the most robust and widely used SG inducer that triggers eIF2α phosphorylation by HRI (10). Exposure to UV light causes eIF2α phosphorylation by a different kinase, GCN2 (12). However, previous studies using eIF2α-S51A mutant human HAP1 cells or MEF[eIF2α-S51A] cells demonstrated that UV-induced SG formation is phospho-eIF2α independent (16, 27). Therefore, we tested if UV inhibits translation initiation and induces SG formation by causing inhibition of mTOR and disassembly of eIF4F complex. Our analysis of the phosphorylation status of mTOR targets 4E-BP and 6SK by western blotting and the assembly of eIF4F complex using m7GTP agarose pulldown assay at 2 h post-UV exposure revealed that mTOR activity and eIF4F complex formation remain unaffected (Fig. 1A,D). When we treated cells with mTOR inhibitor Torin-1, it caused protein synthesis inhibition similar to that which was induced by exposure to 10 mJ/cm^2^ UV, and when combined, the two treatments had an additive effect on the magnitude of translation arrest, indicating that their mechanism of action may be different (Fig. 1B). Importantly, Torin-1 treatment did not enhance SG formation triggered by UV light, providing additional evidence that UV-induced SG formation is not driven by mTOR inhibition (Fig. 1C).

Torin-1 directly inhibits the catalytic activity of mTOR and does not activate ISR or induce SG formation in the absence of stress, even though it causes translation inhibition and polysome disassembly (32). By contrast, amino acid starvation simultaneously causes the inhibition of mTOR activity and accumulation of uncharged tRNAs that activate GCN2 and trigger GCN2-mediated eIF2α phosphorylation (11, 34). SG formation can be induced in HeLa cells following 2 h of amino acid starvation (42). However, similarly to Torin-1 treatment, amino acid deprivation alone did not trigger SG formation in U2OS cells in our system (Fig. 1C). This may be due to differences in response to amino acid starvation between HeLa and U2OS cells or small variations in the exact composition of amino acid free media between our study and the study by Damgaard and Lykke-Andersen (42), nevertheless our results clearly show that translation arrest triggered by amino acid starvation and UV light have different mechanisms despite causing eIF2α phosphorylation by the same kinase GCN2. At later times post-UV exposure, mTOR inhibition and induction of cytoprotective autophagy is shown to be important for cell survival following UV damage (43). However, our data reveals that immediately following UV exposure, translation arrest and SG formation are not caused by mTOR inhibition.

Previous studies have shown that eIF2α phosphorylation is not required for UV-induced SG formation. At the same time, cells engineered to express unphosphorylatable S51A eIF2α mutant are more susceptible to UV damage-induced cell death (16, 41). Therefore, we quantified the contribution of eIF2α phosphorylation to the magnitude of translation arrest following UV exposure. In MEFs expressing a non-phosphorylatable eIF2α (MEF[eIF2α-S51A]), UV treatment inhibited translation, however the magnitude of UV-induced translation arrest measured by ribopuromycylation assay was lower compared to the wild type MEFs (Fig. 2A-C). As expected, translation arrest in MEF[eIF2α-S51A] was not affected by treatment with ISRIB which acts by negating effects of eIF2α phosphorylation on translation initiation. By contrast, ISRIB significantly increased translation rates in both the UV-treated and the control As-treated wild-type MEFs (Fig. 2C). Our results indicate that even though the UV-induced translation arrest mechanism does not act exclusively through eIF2α phosphorylation, it does contribute to its magnitude. At the same time, we confirmed previous reports that UV-induced SG formation is independent of eIF2α phosphorylation (Fig. 2D) (16, 27). Apart from inhibiting bulk protein synthesis, eIF2α phosphorylation allows for preferential synthesis of proteins involved in ISR (15). The best-known example of mRNAs that become translated more efficiently is the one that encodes activating transcription factor 4 (ATF4). In its 5’ region it contains short upstream open reading frames (uORFs) that inhibit initiation of protein synthesis from the downstream ATF4 ORF under normal conditions. When initiation is inhibited by stress-induced eIF2α phosphorylation, the downstream ORF becomes more accessible and expression of ATF4 is induced (44). This transcription factor then translocates to the nucleus where it activates the stress response gene expression program (45). Given that eIF2α phosphorylation is not required for UV-induced translation arrest and SG formation yet plays an important role in cell survival, it is tempting to speculate that its role in regulating synthesis of ISR genes like ATF4 is contributing to a better stress response. However, it was demonstrated previously that due to inhibition of transcription caused by DNA damage, UV exposure does not upregulate ATF4 protein levels (46). Therefore, the exact reason for decreased survival of cells expressing unphosphorylatable eIF2α remains unknown.

Having examined the contribution of eIF2α phosphorylation to the magnitude of translation arrest in response to UV damage, next we examined the role of the eIF2α kinase GCN2. In addition to UV light, this kinase can be activated by other stress stimuli, including amino acid starvation, inhibition of the ubiquitin proteasome system by MG132 treatment, and even viral infections (11, 12, 47, 48). The mechanism of GCN2 activation is best understood for amino acid starvation that leads to accumulation of deacylated tRNAs. GCN2 possesses a histidyl-tRNA synthetase-like (HisRS) domain close to the c-terminus that is involved in autoinhibitory interactions in the inactive GCN2 dimer. Binding of “uncharged” deacylated tRNAs by HisRS domain causes structural rearrangements in the GCN2 dimer interface allowing for autophosphorylation and activation of the kinase (11, 49). Another, tRNA-independent mechanism of GCN2 activation by the P-stalk of ribosomes stalled during elongation have been described, which can complement tRNA-dependent activation when amino acids are depleted (33, 50). How GCN2 is activated by UV light and whether it involves ribosomal stalling on UV-damaged mRNAs is not known, but regardless of triggering stimuli, the activated kinase’s main target is the eIF2α. How many other GCN2 phosphorylation targets exist is presently unknown, however it was shown that it can phosphorylate at least one additional protein: methionyl-tRNA synthetase (MRS) (51). The GCN2-mediated phosphorylation of MRS is directly involved in UV damage response because it causes dissociation of MRS and the aminoacyl-tRNA synthetase-interacting multifunctional protein-3 (AIMP3). The latter then translocates to the nucleus where it participates in DNA damage response. Another consequence of MRS phosphorylation is the inhibition of methionyl-tRNA generation and depletion of the eIF2-GTP-Met-tRNA^Met^ ternary complex that is required for translation initiation, but through the mechanism independent of eIF2α phosphorylation (51). This could potentially explain how UV-induced translation arrest can be largely eIF2α-independent but still require GCN2. However, when we silenced GCN2 expression in U2OS cells using siRNAs, the translation arrest after UV exposure was greatly reduced compared to control cells but still evident (Fig. 3B,C). Importantly, in the presence of ISRIB, the magnitude of translation arrest in cells with silenced GCN2 expression was similar (Fig. 3C). These results, together with MEF data (Fig. 2C), further support the mechanism in which the GCN2-mediated eIF2α phosphorylation contributes significantly to the magnitude of UV-induced translation arrest, but there is little to eIF2α-independent effect of GCN2. By contrast, silencing of GCN2 significantly decreased UV-induced SG formation (Fig. 3D,E), indicating that that it may be involved, directly or indirectly, in the mechanism of SG nucleation. Further analysis using specific inhibitor of GCN2, A-92, showed that the catalytic activity of the GCN2 contributes to UV-induced SG formation mechanism (Fig. 4). Thus, the catalytic activity of GCN2 is important for UV-induced SG formation, but not because of the eIF2α phosphorylation or other direct effects on translation.

Our analysis of UV-induced SG formation revealed that these granules differ from both canonical SGs that require eIF2α phosphorylation and the 4E-BP-dependent SGs that form upon stress-induced inhibition of mTOR. Next, we tested if UV-induced SG condensation requires G3BP1/2 proteins. Depending on the type of stress, G3BP1 and G3BP2 that share 70% amino acid sequence homology can only partially substitute for each other in the mechanism of SG formation (35). In U2OS cells, silencing of either protein causes compensatory increases in the levels of the other, however, disruption of G3BP1 or G3BP2 expression individually still decreased SG formation induced by sodium arsenite. By contrast, formation of SGs induced by clotrimazole treatment is impaired only when both G3BP genes are silenced simultaneously, while osmotic or heat shock cause SG formation by the mechanism that is independent of both G3BP1 and G3BP2 (35). In our study we used CRISPR/Cas9 mediated disruption of G3BP1 or G3BP2 genes in U2OS cells independently and compared contribution of these proteins to UV-induced SG formation. In parallel, we tested the effects of G3BP1 and G3BP2 on SG induction by sodium arsenite and sodium selenite. In our system, disruption of G3BP1 expression prevented UV-induced SG formation but had no effect on SG formation triggered by either arsenite or selenite, as visualised by co-staining with SG markers FMRP and TIA-1 (Fig. 5A). By contrast, both UV and arsenite induced SGs in cells that had G3BP2 expression disrupted by CRISPR/Cas9, however G3BP2 was required for selenite-induced SG formation. Future studies should reveal what differences between G3BP1 and G3BP2 are responsible for selective requirements of these proteins for UV-induced vs. selenite-induced SG formation in U2OS cells, but the reliance on G3BP1 suggests that UV-induced SGs can form via the previously established mechanism that requires binding of Caprin-1 protein to the NTF2 domain of G3BP1 (35). This mechanism of SG formation can be inhibited by ectopic overexpression of the N-terminal peptide of USP10 that contains conserved FGDF sequence motif. This motif is responsible for competitive binding of USP10 to the NTF2 domain of G3BP1 and displacement of Caprin-1 (35). Similarly, several viral proteins that contain FGDF motif can inhibit SG formation in infected cells (52). Consistent with the Caprin-1 dependent mechanism of SG formation, when we overexpressed the N-terminal 40-amino acid peptide of USP10 fused to the EGFP in U2OS cells, UV-induced SG formation was blocked (Fig. 5D). These results convincingly show that untranslated mRNPs that form UV-induced SGs phase separate via the mechanism similar to that of canonical sodium arsenite-induced SGs and requires G3BP1 NTF2 domain interactions.

Having determined that UV-induced SG formation in our system is driven by G3BP1 and is not associated with disassembly of the eIF4F complex, we wanted to confirm that SGs that we detect in U2OS cells at 2 h post-exposure to UV light do not accumulate eIF4G and eIF3B, as was reported by others in different cell types (16, 27). As shown in Fig. 6, unlike SGs that formed in response to sodium arsenite, UV-induced SGs did not recruit eIF4G or eIF3B in our system. Interestingly, while we could detect eIF3B in some SGs induced by the H_2_O_2_ or sodium selenite, even the largest foci in UV-treated cells lacked eIF3B signal (Fig. 6B). To rule out the possibility that UV causes selective dissociation of eIF3 complex from eIF4F, we performed co-immunoprecipitation using eIF3B-specific antibody and analysed eIF3B-eIF4G interaction (Fig. 6C). Similar to sodium arsenite treatment, UV exposure did not disrupt eIF3B-eIF4G interaction. Together with our analysis of eIF4F complex formation using m^7^GTP-agarose pulldown (Fig. 1D), these results indicate that UV-induced translation arrest acts downstream of 48S complex assembly, partially through GCN2-mediated eIF2α phosphorylation and partially through a yet to be identified GCN2 and eIF2α independent mechanism. At the same time, UV-induced SG formation mechanism acts on a subset of mRNPs that lose their association with eIF4G and eIF3 as an additional step downstream of the eIF2α-independent translation arrest (Fig. 6D).

## ACKNOWLEDGEMENTS

We thank Dr. Randal Kaufman (Sanford Burnham Prebys Medical Discovery Institute, La Jolla, CA, USA) and Drs. Adrienne Weeks and Kathleen Attwood (Dalhousie University, Halifax, NS, Canada) for generously providing reagents used in this study. We also thank Dr. Eric Pringle (Dalhousie University, Halifax, NS, Canada) for his help with pulldown assays.

## FUNDING

This work was supported by the Natural Sciences and Engineering Research Council of Canada [grant RGPIN-2019-04323].

## COMPETING INTERESTS

None declared.

## REFERENCES

1. Protter, D.S.W. and Parker, R. (2016) Principles and Properties of Stress Granules. Trends in Cell Biology, 26, 668–679.

2. Kedersha, N., Ivanov, P. and Anderson, P. (2013) Stress granules and cell signaling: more than just a passing phase? Trends in Biochemical Sciences, 38, 494–506.

3. Lin, Y., Protter, D.S.W., Rosen, M.K. and Parker, R. (2015) Formation and Maturation of Phase-Separated Liquid Droplets by RNA-Binding Proteins. Mol. Cell, 60, 208–219.

4. McCormick, C. and Khaperskyy, D.A. (2017) Translation inhibition and stress granules in the antiviral immune response. Nature Reviews Immunology, 17, 647–660.

5. Li, Y.R., King, O.D., Shorter, J. and Gitler, A.D. (2013) Stress granules as crucibles of ALS pathogenesis. J. Cell Biol., 201, 361–372.

6. Baradaran-Heravi, Y., Van Broeckhoven, C. and van der Zee, J. (2020) Stress granule mediated protein aggregation and underlying gene defects in the FTD-ALS spectrum. Neurobiol. Dis., 134, 104639.

7. Monahan, Z., Shewmaker, F. and Pandey, U.B. (2016) Stress granules at the intersection of autophagy and ALS. Brain Res., 1649, 189–200.

8. Kedersha, N., Chen, S., Gilks, N., Li, W., Miller, I.J., Stahl, J. and Anderson, P. (2002) Evidence that ternary complex (eIF2-GTP-tRNA(i)(Met))-deficient preinitiation complexes are core constituents of mammalian stress granules. Mol. Biol. Cell, 13, 195–210.

9. Anderson, P. and Kedersha, N. (2002) Visibly stressed: the role of eIF2, TIA-1, and stress granules in protein translation. Cell Stress Chaperones, 7, 213–221.

10. McEwen, E., Kedersha, N., Song, B., Scheuner, D., Gilks, N., Han, A., Chen, J.-J., Anderson, P. and Kaufman, R.J. (2005) Heme-regulated Inhibitor Kinase-mediated Phosphorylation of Eukaryotic Translation Initiation Factor 2 Inhibits Translation, Induces Stress Granule Formation, and Mediates Survival upon Arsenite Exposure. J. Biol. Chem., 280, 16925–16933.

11. Wek, S.A., Zhu, S. and Wek, R.C. (1995) The histidyl-tRNA synthetase-related sequence in the eIF-2 alpha protein kinase GCN2 interacts with tRNA and is required for activation in response to starvation for different amino acids. Mol. Cell. Biol., 15, 4497–4506.

12. Deng, J., Harding, H.P., Raught, B., Gingras, A.-C., Berlanga, J.J., Scheuner, D., Kaufman, R.J., Ron, D. and Sonenberg, N. (2002) Activation of GCN2 in UV-irradiated cells inhibits translation. Curr. Biol., 12, 1279–1286.

13. García, M.A., Meurs, E.F. and Esteban, M. (2007) The dsRNA protein kinase PKR: virus and cell control. Biochimie, 89, 799–811.

14. Harding, H.P., Zhang, Y., Bertolotti, A., Zeng, H. and Ron, D. (2000) Perk is essential for translational regulation and cell survival during the unfolded protein response. Mol. Cell, 5, 897–904.

15. Jackson, R.J., Hellen, C.U.T. and Pestova, T.V. (2010) The mechanism of eukaryotic translation initiation and principles of its regulation. Nature Reviews Molecular Cell Biology, 11, 113–127.

16. Aulas, A., Fay, M.M., Lyons, S.M., Achorn, C.A., Kedersha, N., Anderson, P. and Ivanov, P. (2017) Stress-specific differences in assembly and composition of stress granules and related foci. J. Cell. Sci., 130, 927–937.

17. Scheuner, D., Song, B., McEwen, E., Liu, C., Laybutt, R., Gillespie, P., Saunders, T., Bonner-Weir, S. and Kaufman, R.J. (2001) Translational Control Is Required for the Unfolded Protein Response and In Vivo Glucose Homeostasis. Molecular Cell, 7, 1165–1176.

18. Sidrauski, C., Acosta-Alvear, D., Khoutorsky, A., Vedantham, P., Hearn, B.R., Li, H., Gamache, K., Gallagher, C.M., Ang, K.K.-H., Wilson, C., et al. (2013) Pharmacological brake-release of mRNA translation enhances cognitive memory. Elife, 2, e00498.

19. Sidrauski, C., McGeachy, A.M., Ingolia, N.T. and Walter, P. (2015) The small molecule ISRIB reverses the effects of eIF2α phosphorylation on translation and stress granule assembly. Elife, 4.

20. Emara, M.M., Fujimura, K., Sciaranghella, D., Ivanova, V., Ivanov, P. and Anderson, P. (2012) Hydrogen peroxide induces stress granule formation independent of eIF2α phosphorylation. Biochemical and Biophysical Research Communications, 423, 763–769.

21. Fujimura, K., Sasaki, A.T. and Anderson, P. (2012) Selenite targets eIF4E-binding protein-1 to inhibit translation initiation and induce the assembly of non-canonical stress granules. Nucleic Acids Research, 40, 8099–8110.

22. Brunn, G.J., Hudson, C.C., Sekulić, A., Williams, J.M., Hosoi, H., Houghton, P.J., Lawrence, J.C. and Abraham, R.T. (1997) Phosphorylation of the translational repressor PHAS-I by the mammalian target of rapamycin. Science, 277, 99–101.

23. Sun, R., Cheng, E., Velásquez, C., Chang, Y. and Moore, P.S. (2019) Mitosis-related phosphorylation of the eukaryotic translation suppressor 4E-BP1 and its interaction with eukaryotic translation initiation factor 4E (eIF4E). Journal of Biological Chemistry, 10.1074/jbc.RA119.008512.

24. Thoreen, C.C., Chantranupong, L., Keys, H.R., Wang, T., Gray, N.S. and Sabatini, D.M. (2012) A unifying model for mTORC1-mediated regulation of mRNA translation. Nature, 485, 109–113.

25. Cadet, J. and Douki, T. (2018) Formation of UV-induced DNA damage contributing to skin cancer development. Photochem. Photobiol. Sci., 17, 1816–1841.

26. Jhappan, C., Noonan, F.P. and Merlino, G. (2003) Ultraviolet radiation and cutaneous malignant melanoma. Oncogene, 22, 3099–3112.

27. Moutaoufik, M.T., El Fatimy, R., Nassour, H., Gareau, C., Lang, J., Tanguay, R.M., Mazroui, R. and Khandjian, E.W. (2014) UVC-Induced Stress Granules in Mammalian Cells. PLoS ONE, 9, e112742.

28. Khaperskyy, D.A., Schmaling, S., Larkins-Ford, J., McCormick, C. and Gaglia, M.M. (2016) Selective Degradation of Host RNA Polymerase II Transcripts by Influenza A Virus PA-X Host Shutoff Protein. PLOS Pathogens, 12, e1005427.

29. Kedersha, N. and Anderson, P. (2007) Mammalian Stress Granules and Processing Bodies. In Methods in Enzymology. Elsevier, Vol. 431, pp. 61–81.

30. Panas, M.D., Kedersha, N. and McInerney, G.M. (2015) Methods for the characterization of stress granules in virus infected cells. Methods, 90, 57–64.

31. Khaperskyy, D.A., Emara, M.M., Johnston, B.P., Anderson, P., Hatchette, T.F. and McCormick, C. (2014) Influenza A Virus Host Shutoff Disables Antiviral Stress-Induced Translation Arrest. PLoS Pathogens, 10, e1004217.

32. Thoreen, C.C., Kang, S.A., Chang, J.W., Liu, Q., Zhang, J., Gao, Y., Reichling, L.J., Sim, T., Sabatini, D.M. and Gray, N.S. (2009) An ATP-competitive mammalian target of rapamycin inhibitor reveals rapamycin-resistant functions of mTORC1. J. Biol. Chem., 284, 8023–8032.

33. Harding, H.P., Ordonez, A., Allen, F., Parts, L., Inglis, A.J., Williams, R.L. and Ron, D. (2019) The ribosomal P-stalk couples amino acid starvation to GCN2 activation in mammalian cells. Elife, 8.

34. Sancak, Y., Peterson, T.R., Shaul, Y.D., Lindquist, R.A., Thoreen, C.C., Bar-Peled, L. and Sabatini, D.M. (2008) The Rag GTPases bind raptor and mediate amino acid signaling to mTORC1. Science, 320, 1496–1501.

35. Kedersha, N., Panas, M.D., Achorn, C.A., Lyons, S., Tisdale, S., Hickman, T., Thomas, M., Lieberman, J., McInerney, G.M., Ivanov, P., et al. (2016) G3BP–Caprin1–USP10 complexes mediate stress granule condensation and associate with 40S subunits. The Journal of Cell Biology, 212, 845–860.

36. Solomon, S., Xu, Y., Wang, B., David, M.D., Schubert, P., Kennedy, D. and Schrader, J.W. (2007) Distinct Structural Features ofCaprin-1 Mediate Its Interaction with G3BP-1 and Its Induction of Phosphorylation of Eukaryotic Translation Initiation Factor 2α, Entry to Cytoplasmic Stress Granules, and Selective Interaction with a Subset of mRNAs. Molecular and Cellular Biology, 27, 2324–2342.

37. Kim, W.J., Back, S.H., Kim, V., Ryu, I. and Jang, S.K. (2005) Sequestration of TRAF2 into stress granules interrupts tumor necrosis factor signaling under stress conditions. Mol. Cell. Biol., 25, 2450–2462.

38. Aulas, A., Lyons, S.M., Fay, M.M., Anderson, P. and Ivanov, P. (2018) Nitric oxide triggers the assembly of “type II” stress granules linked to decreased cell viability. Cell Death & Disease, 9.

39. Reineke, L.C. and Lloyd, R.E. (2015) The Stress Granule Protein G3BP1 Recruits Protein Kinase R To Promote Multiple Innate Immune Antiviral Responses. Journal of Virology, 89, 2575–2589.

40. Batista, L.F.Z., Kaina, B., Meneghini, R. and Menck, C.F.M. (2009) How DNA lesions are turned into powerful killing structures: Insights from UV-induced apoptosis. Mutation Research/Reviews in Mutation Research, 681, 197–208.

41. Burgess, H.M., Richardson, W.A., Anderson, R.C., Salaun, C., Graham, S.V. and Gray, N.K. (2011) Nuclear relocalisation of cytoplasmic poly(A)-binding proteins PABP1 and PABP4 in response to UV irradiation reveals mRNA-dependent export of metazoan PABPs. J Cell Sci, 124, 3344–3355.

42. Damgaard, C.K. and Lykke-Andersen, J. (2011) Translational coregulation of 5’TOP mRNAs by TIA-1 and TIAR. Genes & Development, 25, 2057–2068.

43. Chen, L.-H., Chu, P.-M., Lee, Y.-J., Tu, P.-H., Chi, C.-W., Lee, H.-C. and Chiou, S.-H. (2012) Targeting Protective Autophagy Exacerbates UV-Triggered Apoptotic Cell Death. Int J Mol Sci, 13, 1209–1224.

44. Harding, H.P., Novoa, I., Zhang, Y., Zeng, H., Wek, R., Schapira, M. and Ron, D. (2000) Regulated translation initiation controls stress-induced gene expression in mammalian cells. Mol. Cell, 6, 1099–1108.

45. Harding, H.P., Zhang, Y., Zeng, H., Novoa, I., Lu, P.D., Calfon, M., Sadri, N., Yun, C., Popko, B., Paules, R., et al. (2003) An integrated stress response regulates amino acid metabolism and resistance to oxidative stress. Mol. Cell, 11, 619–633.

46. Dey, S., Baird, T.D., Zhou, D., Palam, L.R., Spandau, D.F. and Wek, R.C. (2010) Both transcriptional regulation and translational control of ATF4 are central to the integrated stress response. J. Biol. Chem., 285, 33165–33174.

47. Mazroui, R., Di Marco, S., Kaufman, R.J. and Gallouzi, I.-E. (2007) Inhibition of the ubiquitin-proteasome system induces stress granule formation. Mol. Biol. Cell, 18, 2603–2618.

48. Berlanga, J.J., Ventoso, I., Harding, H.P., Deng, J., Ron, D., Sonenberg, N., Carrasco, L. and de Haro, C. (2006) Antiviral effect of the mammalian translation initiation factor 2alpha kinase GCN2 against RNA viruses. EMBO J., 25, 1730–1740.

49. Masson, G.R. (2019) Towards a model of GCN2 activation. Biochem. Soc. Trans., 10.1042/BST20190331.

50. Inglis, A.J., Masson, G.R., Shao, S., Perisic, O., McLaughlin, S.H., Hegde, R.S. and Williams, R.L. (2019) Activation of GCN2 by the ribosomal P-stalk. Proc. Natl. Acad. Sci. U.S.A., 116, 4946–4954.

51. Kwon, N.H., Kang, T., Lee, J.Y., Kim, H.H., Kim, H.R., Hong, J., Oh, Y.S., Han, J.M., Ku, M.J., Lee, S.Y., et al. (2011) Dual role of methionyl-tRNA synthetase in the regulation of translation and tumor suppressor activity of aminoacyl-tRNA synthetase-interacting multifunctional protein-3. Proc. Natl. Acad. Sci. U.S.A., 108, 19635–19640.

52. Panas, M.D., Schulte, T., Thaa, B., Sandalova, T., Kedersha, N., Achour, A. and McInerney, G.M. (2015) Viral and cellular proteins containing FGDF motifs bind G3BP to block stress granule formation. PLoS Pathog., 11, e1004659.

